# Subfoveal scotomas trigger fine-scale fixation reorganization: insights from retinal imaging and retinal-contingent stimulation

**DOI:** 10.1101/2025.09.17.674062

**Authors:** Benjamin Moon, Ashley M. Clark, Krishnamachari S. Prahalad, Austin Roorda, Pavan Tiruveedhula, Wolf Harmening, Aleksandr Gutnikov, Samantha K. Jenks, Sanjana Kapisthalam, Michele Rucci, Jannick P. Rolland, Martina Poletti

## Abstract

Fine spatial vision relies on the foveola, the 1-degree retinal region with highest cone density. Despite its importance, the relationship between retinal anatomy, fixational behavior, and visual perception in the foveola is not fully understood. Using an Adaptive Optics Scanning Light Ophthalmoscope for high-resolution retinal imaging and stimulation, we studied the effect of a simulated subfoveolar (*≈*0.03 degrees^2^) scotoma on fine spatial vision and fixation behavior in healthy observers. Our findings show that the visuomotor system adapts to the scotoma with striking precision by shifting the preferred locus of fixation in a systematic fashion by minute (*≈*5 arcmin) amounts to bring stimuli into a region of visibility. These results reveal an unprecedented level of fine-scale plasticity in the human visuomotor system. Interestingly, this new retinal locus of fixation is characterized by lower cone density among those surrounding the scotoma, indicating that factors beyond spatial sampling maximization influence these fine-scale adjustments.

## Introduction

Vision is not uniform across the visual field; it is well known that visual resolution and other functions gradually decrease as we move away from the center of gaze. High-acuity vision is limited to the central fovea, also referred to as the foveola, a morphologically distinct region from the rest of the retina which spans only one degree of visual angle, about the size of an index fingernail at arm’s length (Tuten and Harmening, 2021, Kolb et al.). The foveola sits in a pit on the retina, is free from rods and capillaries, and is characterized by the highest cone packing. Cone density is not uniform in this small region (Curcio et al., 1990) and it reaches a peak within a small portion of the foveola where cone density can reach even 20,000 cones/degree^2^ (Reiniger et al., 2021, Wang et al., 2019). Central foveal vision is essential for performing high-acuity tasks, and the loss or impairment of foveal vision through retinal degenerative diseases has a profound negative impact on quality of life (Hazel et al., 2000, Sengupta et al., 2015, Chung, 2020).

Interestingly, foveal scotomas not only impact vision but also oculomotor behavior. In diseases such as Age-related Macular Degeneration (AMD), a central scotoma can impact much or all of the fovea, resulting in severely impaired visual acuity. Once vision in the central fovea is lost binocularly, patients adopt an alternative and more eccentric preferred retinal locus (PRL) to compensate for this loss; that is, oculomotor behavior changes and saccades are re-referenced to an alternative PRL (Verghese et al., 2021, White and Bedell, 1990), which is generally located near the scotoma border, as illustrated in Figure 1A (Sunness et al., 1996, Fletcher and Schuchard, 1997, Sunness and Applegate, 2005, Crossland et al., 2005, Vullings and Verghese, 2021). Fixational stability is also drastically impaired in these patients (Bellmann et al., 2004, Krishnan and Bedell, 2018, Chung, 2020). Similarly to what happens in patients, in human observers with normal vision a simulated foveal scotoma not only leads to impaired visual performance, but also affects oculomotor behavior and an alternative PRL develops in a relatively short period of time (Kwon et al., 2013, Liu and Kwon, 2016, Barraza-Bernal et al., 2018, Prahalad and Coates, 2020).

**Figure 1:**
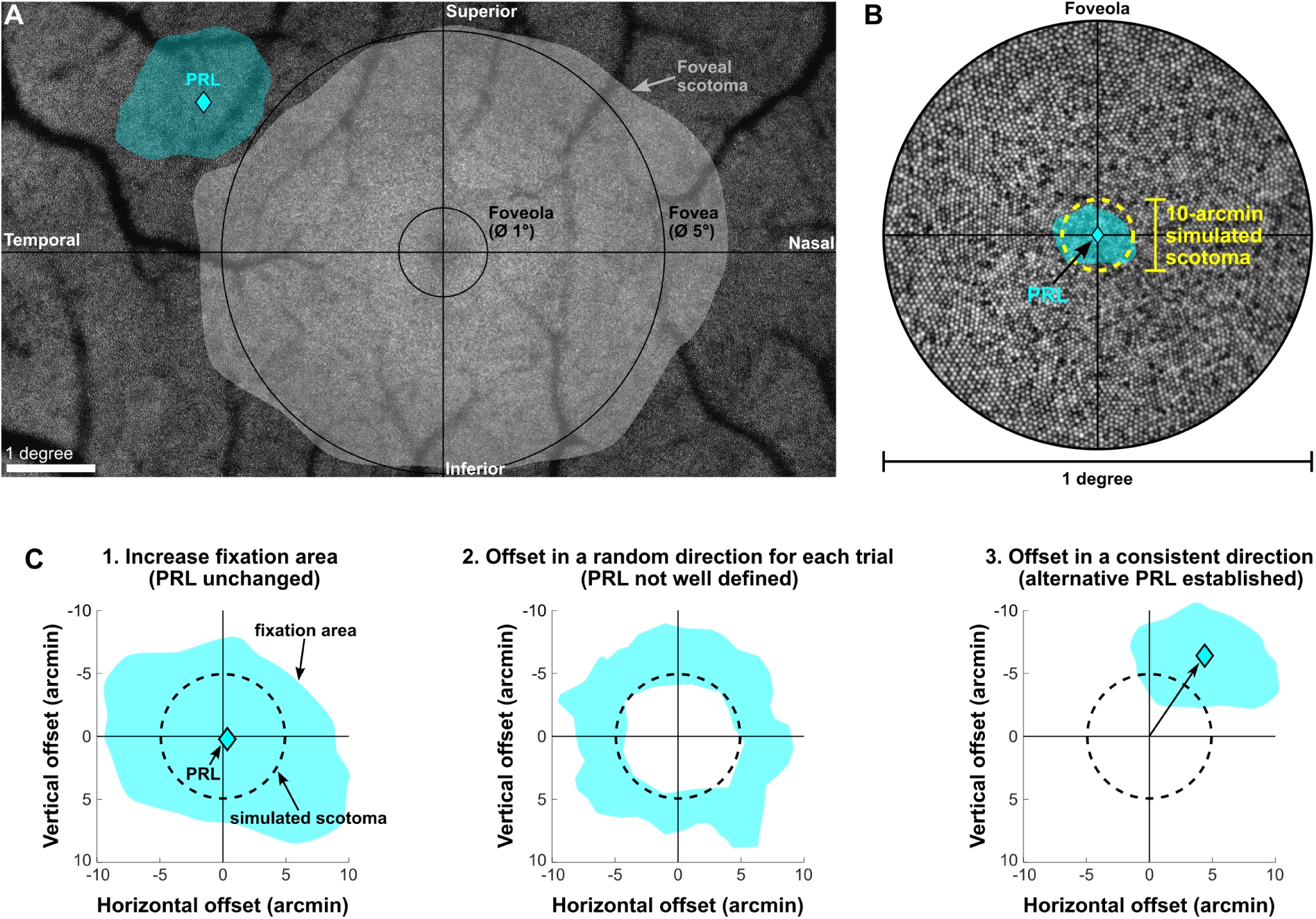
Comparison of foveal and subfoveal scotomas. (A) For scotomas that span the entire fovea, as pictured by the semi-transparent gray region, it is well established that subjects and patients develop an alternative PRL near the scotoma border, as illustrated by the cyan diamond inside the cyan-colored fixation area. The most common alternative PRL location is in the superior-temporal retina (for the right eye pictured here), corresponding to the scotoma being above and to the right of fixation in the visual field (Crossland et al., 2005, Sunness and Applegate, 2005) (B) For subfoveal scotomas with dimensions comparable to the fixation area for an observer with normal fixational stability, it is unknown to what extent oculomotor behavior is influenced by the presence of the small scotoma. In this study, we used a 10-arcmin-wide simulated scotoma that was stabilized at each subject’s PRL, or center of gaze. (C) Three predictions describing possible changes in oculomotor behavior in response to the simulated scotoma. 1. Observers may increase their fixational eye movements to bring the stimulus into a region of visibility without changing the average gaze position, thus maintaining the same PRL. 2. Observers may offset in a random direction for each trial to see the stimulus without establishing a stable alternative PRL. 3. Observers may offset in a consistent direction across trials to establish a stable alternative PRL near the border of the simulated scotoma.

Research has primarily focused on visual and oculomotor effects of large (*>*1 degree) foveal scotomas because in AMD, patients may initially be unaware of their vision loss, which often starts in the more peripheral part of the macula, and their scotomas may span several degrees before diagnosis (Bressler, 2002, Fletcher et al., 2012). Additionally, it is challenging to render subfoveal scotomas in simulations of vision loss using traditional eye-tracking methods because uncertainty in localization of the center of gaze is typically around 1 degree (Holmqvist et al., 2011, Blignaut and Wium, 2014), making it difficult to precisely stabilize a sub-degree scotoma within the foveola (see Figure 1B, which illustrates an example of a small subfoveal scotoma with an area comparable to the 68% fixation contour surrounding the PRL for a vision-healthy observer). Furthermore, incessant involuntary eye movements constantly shift the retinal image over many photoreceptors during fixation (Ratliff and Riggs, 1950, Barlow, 1952, Steinman et al., 1973, Kowler, 2011, Rucci and Poletti, 2015). Therefore, to simulate vision loss from a subfoveal scotoma, eye movements must be measured with high precision and compensated in near-real-time to ensure that the simulated scotoma remains stationary on the retina. Because of these challenges, it has remained unknown to what extent the visual system is sensitive to small subfoveal scotomas and whether oculomotor behavior changes to compensate for such losses.

Quantifying the impact on vision that results from a subfoveal scotoma has far-reaching implications in understanding central foveal vision and its relationship with oculomotor behavior. Vision within the foveola is often assumed to be uniform. However, while sensitivity to small spots of light is fairly uniform within this region (Domdei et al., 2021), neither visual discrimination (Poletti et al., 2013, Intoy and Rucci, 2020) and visual detection in noise (Intoy et al., 2021), nor cone density is constant across the foveola (illustrated in Figure 2A for an example subject; see also Curcio et al. (1990), Rossi and Roorda (2010), Wang et al. (2019), Reiniger et al. (2021)). Further, there are foveal asymmetries in vision-healthy observers that arise from small offsets between the PRL and the point of peak cone density (PCD) and the Cone Density Centroid (CDC). The CDC is defined as the weighted centroid of the top 20% of cone density values, and has been shown to more accurately represent the anatomical center of the fovea compared with the PCD location (Reiniger et al., 2021, Wynne et al., 2022, Warr et al., 2024). It is well known that the preferred retinal locus does not generally align with these anatomical landmarks (Putnam et al., 2005, Li et al., 2010, Wilk et al., 2017, Wang et al., 2019, Kilpeläinen et al., 2021), and there is instead a systematic offset between these locations (Reiniger et al., 2021) (Figure 2B). This offset introduces an asymmetry in cone density along the vertical meridian (see Figure 2C). As a result, the average observer is characterized by a higher cone density in the inferior-temporal retina (upper left foveal visual field for the right eye) relative to the PRL.

**Figure 2:**
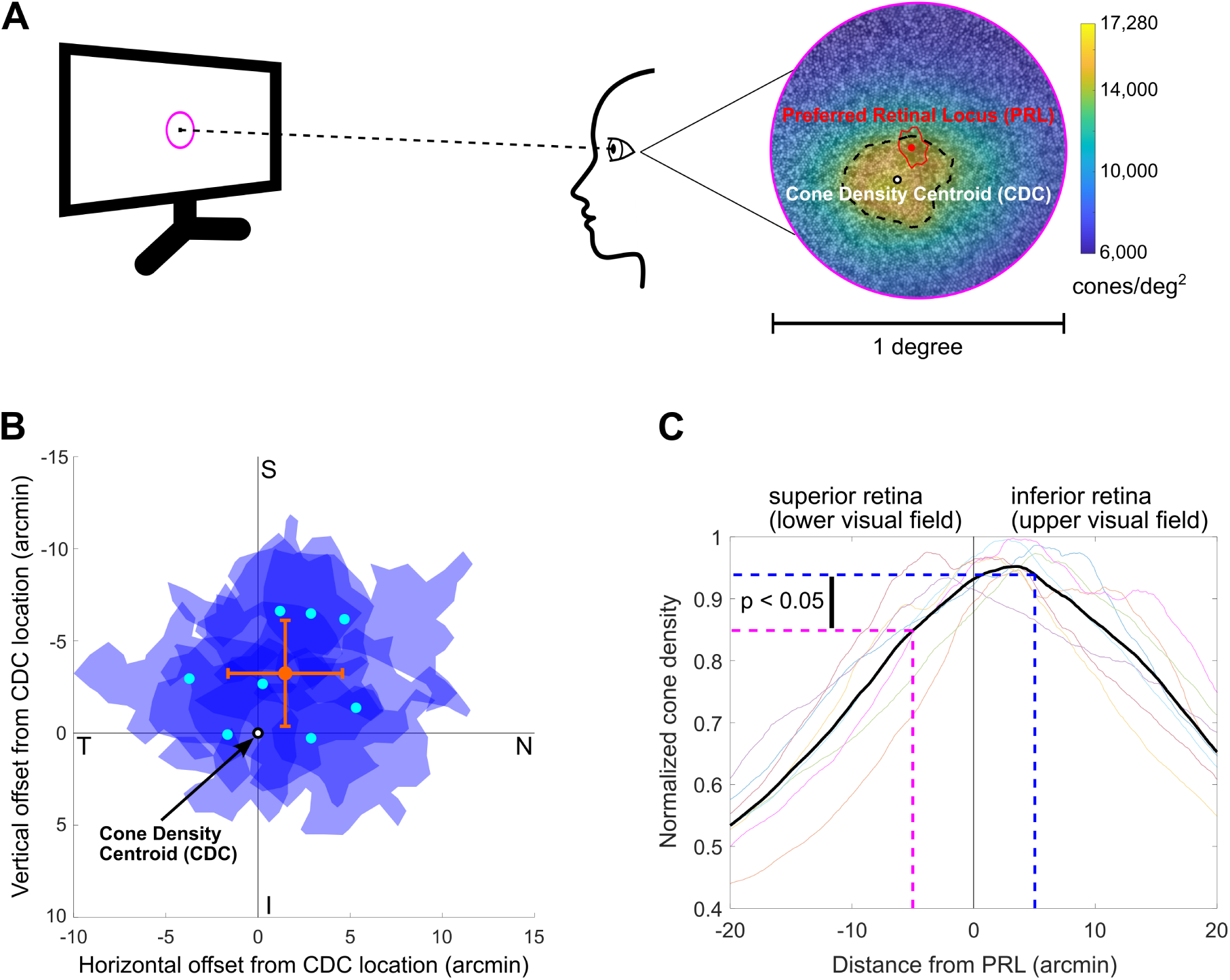
Offset between the cone density centroid and the preferred retinal locus. (A) Human observers with normal vision center stimuli on a specific location within the foveola, called the Preferred Retinal Locus, or PRL. Cone density within the foveola is not uniform, but decreases by approximately a factor of two from the Cone Density Centroid (CDC) to the edge of the 1-degree-diameter foveola. The PRL rarely coincides with the CDC, but is instead offset by a few arcmin. (B) Across eight observers, we found an average PRL offset of 4.9 *±* 2.3 arcmin from the CDC location, with the average offset occurring toward the superior-nasal retina with respect to the CDC. The error bars denote the standard deviations along the horizontal and vertical axes. (C) This offset of the PRL from the CDC creates an asymmetric distribution of photoreceptor sampling across the visual scene, with higher sampling density occurring in the upper visual field for the average observer (t(7) = −2.69, p = 0.03, two-tailed paired t-test).

This retinal and perceptual diversity characterizing the foveolar landscape raises a number of questions on the interplay between fixational behavior and visual perception. Does a small subfoveal scotoma lead to a substantial drop in acuity? Does fixation behavior change in response to this visual alteration? Developing an alternative PRL at this fine scale is not trivial as oculomotor adjustments would be on the order of arcminutes. In addition, it may not even be necessary to develop an alternative PRL to maintain high-acuity vision, as increasing fixational instability may be sufficient to bring the stimulus into a region of visibility (see Figure 1C). On the other hand, if an alternative PRL is established, it becomes important to understand if its formation is influenced by a process maximizing spatial sampling of the visual input, moving it toward the highest cone density region that is accessible outside the scotoma. To address these questions we used a high-resolution Adaptive Optics Scanning Light Ophthalmoscope (AOSLO) to simultaneously image the foveal cone mosaic at high resolution, present stimuli in a retinally contingent manner to render a simulated subfoveal scotoma, and record fixational oculomotor behavior at high resolution. This experimental setup enables maintaining the simulated scotoma over a fixed array of cones throughout the fixation period. Thanks to this sophisticated technology, this study unravels the finesse of fixational oculomotor behavior and the diversity of central foveal vision. Further, it sheds light on the mechanisms driving the establishment of an alternative PRL. It also has important clinical implications highlighting that fine changes in fixation behavior may be associated with the presence of subfoveal vision loss.

## Results

To examine the impact of a subfoveal scotoma on visual performance and fixation behavior, we used an AOSLO to render a simulated 10-arcmin-wide scotoma while subjects (n = 7) performed a four-alternative forced choice (4AFC) visual acuity discrimination task. Stimuli consisted of optotypes whose size (5 arcmin in width, corresponding to 20/20 Snellen acuity) was considerably above the acuity threshold as they were projected on the retina bypassing optical aberrations. Stimuli were presented for 500 ms at a fixed location on the display. The task was either performed in the presence or absence of the scotoma (Figure 3B). In the scotoma condition, the 10-arcmin-wide simulated scotoma, whose borders were invisible to the observers, was stabilized at each subject’s PRL. Each trial was initiated by a button press from the subject, and trials were self-paced. 250 ms before the stimulus onset, a 1-second recording of the retinal cone mosaic was initiated for each trial (Figure 3C). These recordings showed the precise locations of both the optotype and the simulated scotoma on the retina, which made it possible to track the stimulus motion on the retina and verify that the scotoma was well stabilized at each subject’s PRL.

**Figure 3:**
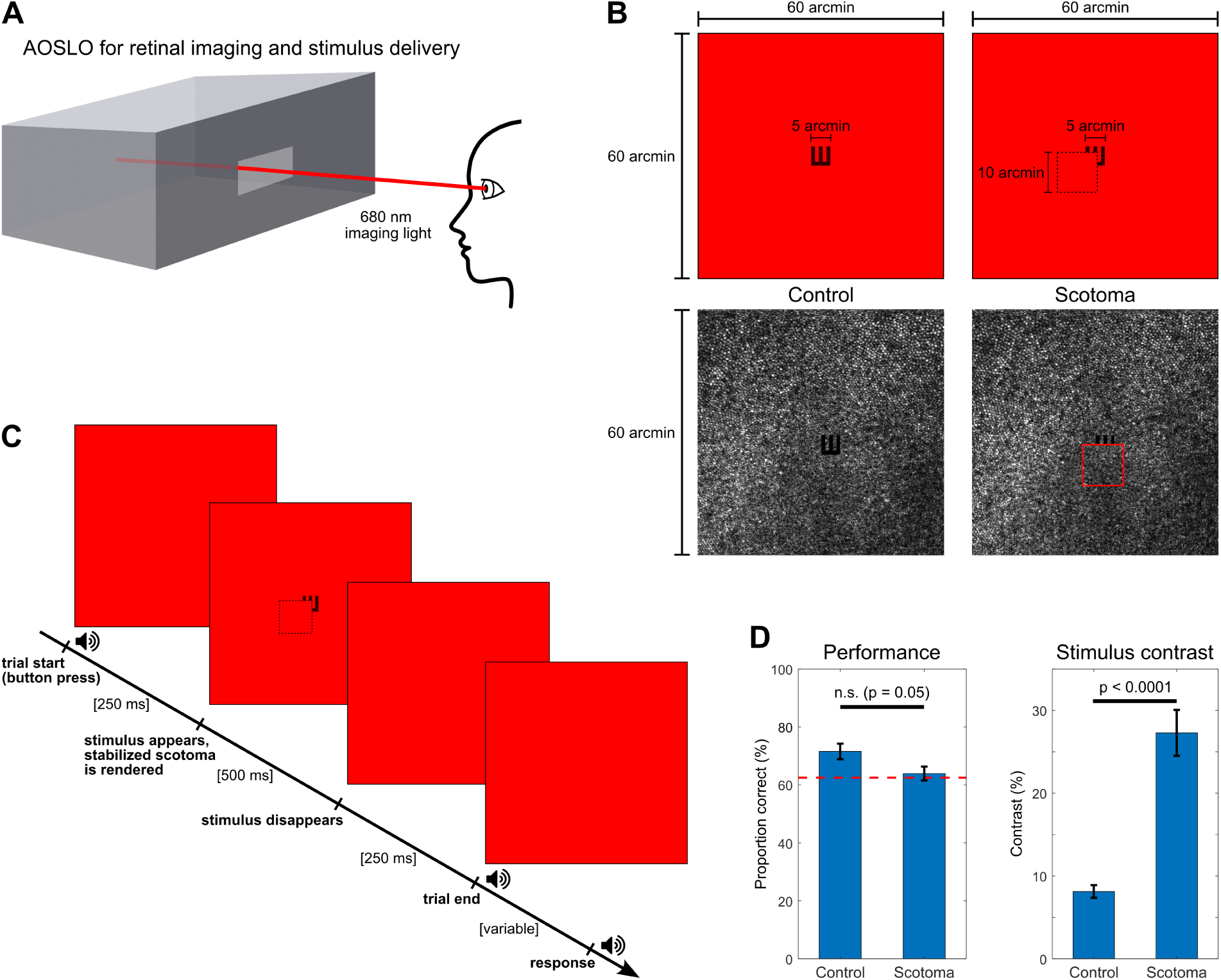
Methods for retinal imaging and retinal-contingent stimulus manipulation. (A) Retinal imaging and stimulus delivery was conducted using a custom Adaptive Optics Scanning Light Ophthalmoscope (AOSLO), which enabled high-resolution imaging of the foveal cones and precise stimulus delivery. All imaging and stimulus delivery utilized 680 nm light, which appeared as a bright red/orange color to observers. (B) Decrement stimuli were presented within the 60-arcmin-wide imaging field of view by modulating the laser light during the raster scan to produce a 5-arcmin-wide Snellen E at the center of the raster scan. The top row shows the appearance of the stimulus to the observer, and the bottom row shows single video frames from a retinal recording of an example observer (note that these video frames have been flipped vertically to match the orientation of the optotypes from the observer’s perspective). In the scotoma condition (column 2), a 10-arcmin-wide simulated scotoma was stabilized at each observer’s PRL, which is shown partially occluding the stimulus in the example video frame, with the scotoma border drawn in red for reference. (C) A 1-second video was recorded for each trial, with the 500-ms stimulus presentation occurring in the middle of the trial. Trials were self-paced, and were initiated by a button press. Control and scotoma blocks utilized the same paradigm, except that the scotoma was not rendered during control blocks. (D) Performance was at or slightly above threshold (denoted by the dashed red line) in both conditions. There was no significant difference in performance between the conditions (t(6) = 2.42, p = 0.052, two-tailed paired t-test). The stimulus contrast was 3.3 *±* 0.3 times higher, on average, in the scotoma condition to achieve similar performance (t(6) = −9.25, p *<*0.0001, two-tailed paired t-test). Error bars denote Standard Error of the Mean: SEM.

The stimulus contrast was adjusted to achieve performance near threshold (*≈* 62% correct responses for our 4AFC task) in both the scotoma and control conditions (Figure 3D). The presence of the scotoma made the task much more difficult; to achieve comparable performance in both conditions, contrast was increased by more than a factor of three on average (3.3 *±* 0.3, t(6) = −9.25, p *<*0.0001, two-tailed paired t-test, see methods for detail) (Figure 3D, right panel) in the scotoma condition. These results show that a small scotoma at the center of gaze, with a width of only 10 arcmin, causes a substantial deficit in performance in a visual acuity task.

We then investigated to what extent oculomotor behavior changed in response to the simulated scotoma. Changing fixation behavior to compensate for the loss of visual information brought by such a small scotoma is not trivial and it would require a high level of finesse and control. A likely possibility would be that fixation behavior remains unchanged and the visual system simply relies on a few glimpses of a high-contrast stimulus to perform the task. In fact, fixation area may be large enough to bring the stimulus into a region of visibility in some trials. Yet, this strategy may not be very effective as most of the time the fixation area is only slightly larger than the region covered by the scotoma. Figure 1C outlines three possible changes in behavior that might be expected in response to the simulated scotoma. First, the visual system could increase its fixational area to bring the stimulus outside of the scotoma borders during a greater proportion of each trial duration. This could be accomplished by increasing the amplitude or speed of ocular drift. In this scenario, the PRL would not change, but the distribution of retinal stimulus coordinates would broaden. Alternatively, the visual system could adopt a random offset in each trial to bring the stimulus into a region of visibility surrounding the scotoma without establishing an alternative PRL. Another possibility is that a consistent offset could be utilized to establish an alternative PRL near the border of the simulated scotoma, similar to what is observed for a large foveal scotoma. The latter scenario in particular would require a remarkably orchestrated effort and coordination between the visual and the motor system.

### Oculomotor behavior during stimulus presentation

To determine whether the simulated scotoma caused a systematic change in oculomotor behavior across subjects, we measured the stimulus motion on the retina due to fixational eye movements during each 500-ms stimulus presentation interval using offline video registration software (Zhang et al., 2021). Fixation behavior is characterized by microsaccades and drift. The microsaccade rate was low in our task (0.15 *±* 0.20 per second in the control condition and 0.23 *±* 0.25 per second in the scotoma condition, with no significant difference between conditions: t(6) = −0.57, p = 0.59, two-tailed paired t-test), so our analysis focuses primarily on drift-only trials. Individual drift segments and drift velocity distributions for an example subject are displayed in Figure 4A and 4B, where the line color denotes the time from stimulus onset. In both the control (top row) and scotoma (bottom row) conditions, this example subject tended to drift in a specific direction, moving the stimulus toward the CDC and the inferior retina. We then compared the mean drift direction with the angle between the PRL and the CDC. As recently reported in Witten et al. (2024), we found that subjects exhibited a high probability of drifting toward their CDC, as shown in Figure 4C, with an average of 66 *±* 12% of the drift velocity distribution moving toward the CDC and 34 *±* 12% moving away from the CDC in the control condition (ANOVA F(1,6) = 15.40, p = 0.008, HSD Test for multiple comparisons p = 0.010 for probability toward vs. away). Interestingly, this trend was less pronounced when subjects did not perform an acuity task and were simply maintaining fixation on a marker of the same size as the high-acuity stimulus (60 *±* 10%, ANOVA F(1,6) = 15.40, p = 0.008, HSD Test for multiple comparisons p = 0.037 for probability toward vs. away), suggesting, in line with recent evidence (Intoy et al., 2024), some degree of active control over this behavior. In the scotoma condition drift behavior was unchanged; subjects demonstrated the same high probability of moving toward the CDC, with 65 *±* 10% of the drift velocity distribution moving toward the CDC. There was no significant difference in any other drift properties (see methods for detail), suggesting that subjects do not systematically change their dominant drift direction in response to a subfoveal scotoma.

**Figure 4:**
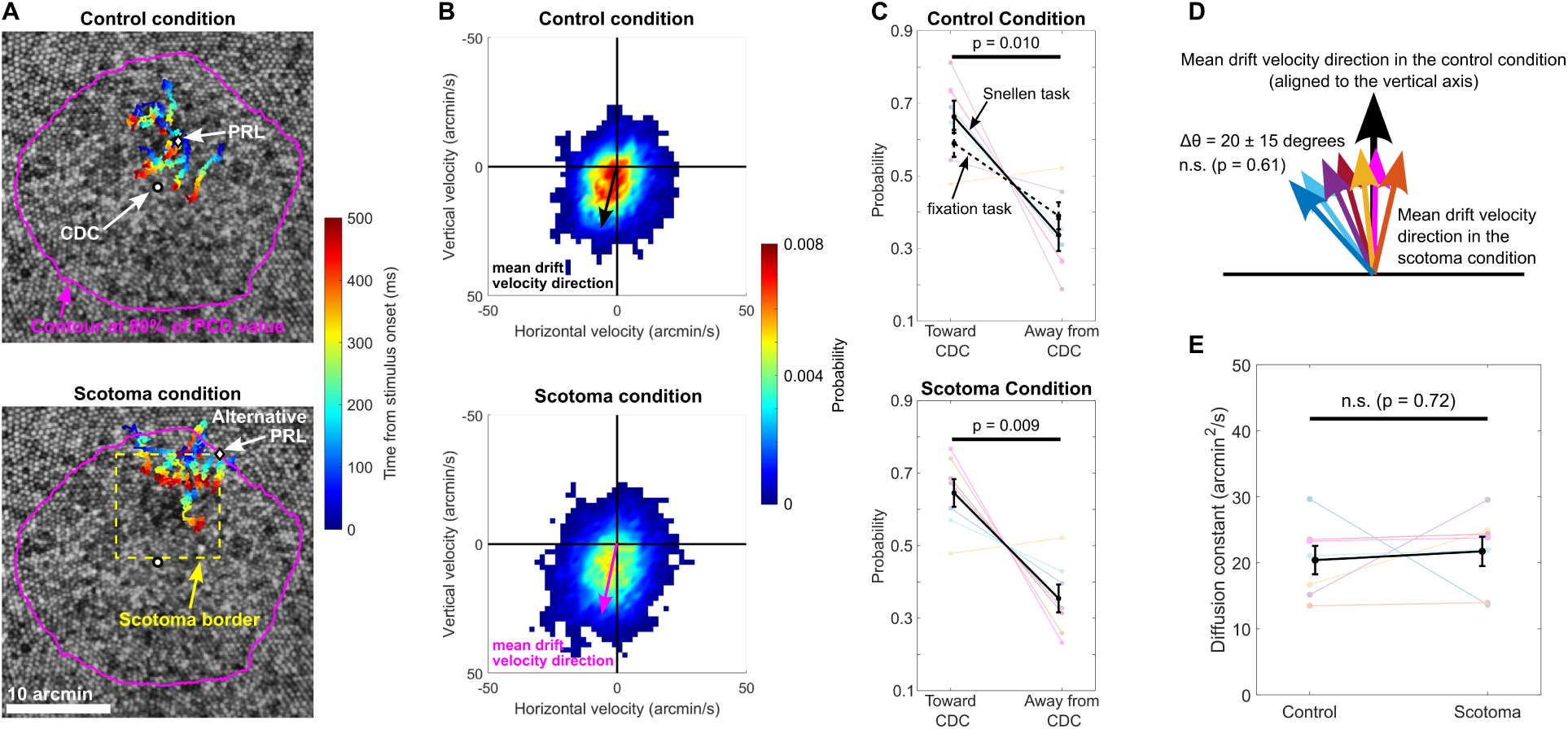
Analysis of drift behavior in the control and scotoma conditions. (A) Individual drift segments are plotted on the cone mosaic for an example subject in both the control (top) and scotoma (bottom) conditions, with the line color representing the time from stimulus onset. Ten example drift segments are plotted. The drift has a directional bias that tends to bring the stimulus closer to the CDC. In the scotoma condition, the drift segments are shifted further from the CDC due to the scotoma occlusion. (B) The directional bias of drift for this example subject is further demonstrated by plotting the distribution of drift velocity across all drift segments in the control (top) and scotoma (bottom) conditions. The black arrow denotes the mean amplitude and direction of drift velocity in the control condition, and the magenta arrow shows the mean amplitude and direction of drift velocity in the scotoma condition. (C) This directional bias of drift velocity was observed across subjects in the control condition (top), as demonstrated by the higher probability of the drift direction moving toward the CDC relative to the PRL. The dashed black line shows the average probability in a separate fixation task, which has a similar trend. In the scotoma condition (bottom), the same average trend is observed, with no significant difference between the control and scotoma conditions (ANOVA F(2,12) *<*0.001, p *>*0.999). Error bars denote SEM. (D) The mean drift direction in the control condition has been aligned to the vertical axis for all subjects, and the colored arrows show the mean velocity direction in the scotoma condition. There was no significant difference in the mean direction of drift velocity between the control and scotoma conditions across subjects (F(1,12) = 0.28, p = 0.61, Watson-Williams test for circular statistics). Δ*θ* describes the mean magnitude of the difference in angle between the control and scotoma mean drift velocity angles. (E) Across subjects, we found no systematic difference in the drift diffusion constant between the control and scotoma conditions (t(6) = −0.38, p = 0.72, two-tailed paired t-test). Error bars denote SEM.

### An alternative PRL is established

So far we examined drift and overall fixation behavior characteristics in the control and in the scotoma condition. Although drift and fixation behavior did not systematically change in the two conditions, it is possible that the retinal location used to center the target on is different. To determine whether an alternative PRL was utilized, we analyzed the distribution of stimulus coordinates on the retina. Our results show that in the scotoma condition all subjects had an offset greater than 2 arcmin between the control PRL and alternative PRL in the scotoma condition (Figure 5A–C). The offsets for each subject, shown by the black arrows in Figure 5C, were not oriented in any systematic direction. Across subjects, the mean offset magnitude between the control PRL and the alternative PRL was 5.5 *±* 2.9 arcmin, and the mean overlap in retinal area encompassed by both the control and scotoma contours was 38 *±* 35%. This finding indicates that the visuomotor system responded to the scotoma by shifting the stimulus distribution away from the PRL by approximately half the width of the scotoma, bringing the stimulus into a region of partial visibility near the scotoma border. What may have caused the shift in the PRL given that the oculomotor behavior during stimulus presentation was comparable in the scotoma and control condition? It is likely that, given the self-paced nature of the task, subjects used microsaccades before the stimulus onset to move their PRL away from the location where the stimulus would appear, so the target would be presented at the desired retinal location during the task.

**Figure 5:**
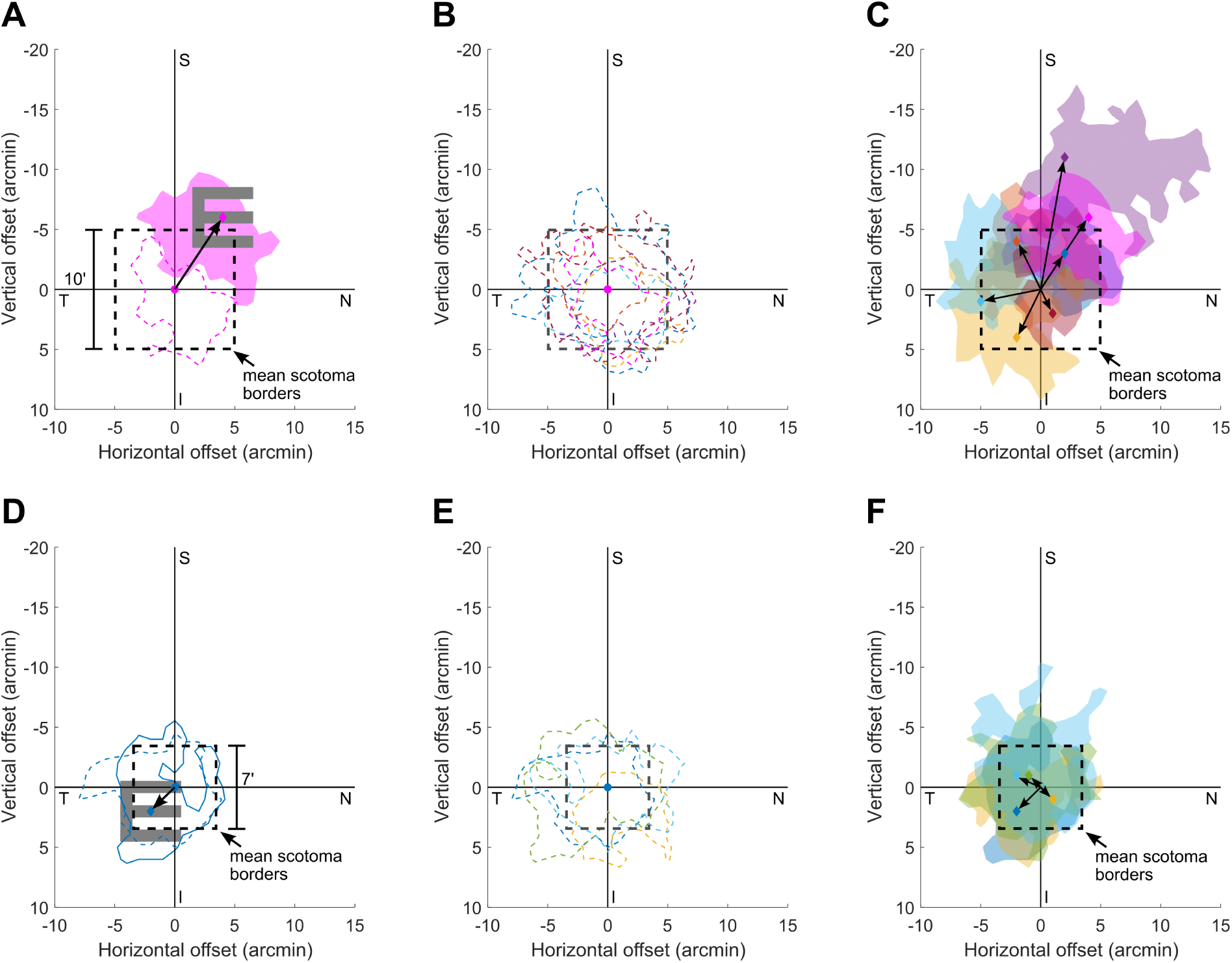
Stimulus distribution offsets, showing the establishment of an alternative PRL for the 10-arcmin-wide scotoma. (A) Offset from the PRL in the scotoma condition for an example subject, with the pink diamond and shaded region denoting the alternative PRL and the dashed contour denoting the control PRL. The offset of 7 arcmin for this example subject brings the E stimulus into a region of visibility outside of the scotoma occlusion (black dashed box). (B) 68% contours for all subjects in the control condition, with the origin centered on each subject’s control PRL. The average area of these contours (63 *±* 29 arcmin^2^), is smaller than the scotoma area of 100 arcmin^2^ (the scotoma borders are shown for reference only: the scotoma was not rendered in the control condition). (C) Contours for all subjects in the scotoma condition. The black arrows show the direction and magnitude of the alternative PRL offsets. The average offset magnitude was 5.5 *±* 2.9 arcmin. (D) When a smaller scotoma with a width of 7 arcmin was tested with 4 subjects, the offsets were considerably smaller. This example subject showed an offset of 2.8 arcmin, which only brings the edge of the E stimulus into the region of visibility, on average. (E) Contours for the control condition for the 4 subjects who participated in the 7-arcmin condition, which span a larger retinal area on average compared with the scotoma area (the scotoma borders are shown for reference only: the scotoma was not rendered in the control condition). (F) Contours for the 4 subjects in the 7-arcmin scotoma condition, with the black arrows denoting the direction and magnitude of the offsets from the control PRL. The average offset magnitude was 2.1 *±* 0.7 arcmin.

Interestingly, in a control experiment in which a smaller (7-arcmin-wide) scotoma was used, no alternative PRL was established. The smaller scotoma, covering only 49 arcmin^2^, occluded a retinal area that was substantially smaller than the average 68% contour area of 64 *±* 19 arcmin^2^ of fixational instability. As a result, normal fixational instability would be sufficient to bring the stimulus into the visible region for a duration long enough to perform the task significantly above chance level even if the stimulus contrast was the same as in the control condition. Contrast level was maintained equal in the two conditions and the average performance in the control condition was 79 *±* 5% correct, and the average performance in the scotoma condition was 66 *±* 9% correct (t(3) = 1.99, p = 0.14, two-tailed paired t-test; Figure 5 D–F). The example subject shown in Figure 5D had an offset of 2.8 arcmin between the control and scotoma conditions, which was the largest observed offset in this control experiment. Figure 5E demonstrates that the 68% contours for the control condition were larger than the scotoma occlusion, which was not the case for the 10-arcmin-wide scotoma. The mean offset in the scotoma condition was 2.1 *±* 0.7 arcmin, which is smaller than the alternative PRL offsets in the 10-arcmin condition (t(9) = 2.29, p = 0.047, unpaired t-test). Furthermore, the 2-arcmin offset is also smaller than half the scotoma width (i.e., 3.5 arcmin). In the 7-arcmin-scotoma experiment, the stimulus was fully occluded in 31 *±* 21% of the video frames. In the 10-arcmin-scotoma experiment, the stimulus was occluded in 55 *±* 16% of the video frames, and there was not a significant difference in the percentage of fully occluded frames between these two experiments (t(9) = −2.20, p = 0.056, unpaired t-test). Altogether, these results suggest that subjects did not establish an alternative PRL for a 7-arcmin-wide scotoma, pointing to the possibility that there exists a minimum occlusion size before substantial changes in fixation behavior are observed.

### The alternative PRL is characterized by a lower cone density compared to the other regions surrounding the scotoma

The offset direction of the alternate PRL in the scotoma condition was not systematic across subjects (Figure 5C), yet the chosen retinal location may have some common anatomical characteristics across subjects. In fact, cone density around the scotoma region was not uniform. If the goal of the visual system is to maximize sampling, we would expect the alternate PRL to be centered on the region of highest cone density available outside the scotoma. Here we investigated to what extent each subject’s offset was related to their unique cone density distribution. We first determined the average cone density at different locations surrounding the scotoma (see methods and Figure 6A and 6B for details). After quantifying the average cone density surrounding the scotoma, we aligned each subject’s polar plot so that the alternative PRL offset direction was oriented vertically, as shown by the black arrow in Figure 6C. The polar plot in Figure 6B shows the average cone density as a function of angle for an example subject. The highest cone density occurred at an angle of 112.5 degrees relative to the center of the scotoma (angles were measured in a clockwise direction to keep all coordinates consistent with the retinal images, which had their origins in the top left corner of the image). The lowest cone density around the scotoma occurred at 315 degrees. For the example subjects shown in Figure 6B, the alternative PRL was offset toward the lowest cone density region surrounding the scotoma. Surprisingly, we found that all subjects set their alternate PRL away from the highest cone density region available outside the scotoma occlusion, and toward a lower cone density region (see Figure 3 in Supplemental Information for individual plots of cone density surrounding the scotoma in retinal coordinates). The angles corresponding to each subject’s highest cone density region are denoted by the triangles in Figure 6C, and all the triangles are below the horizontal axis, indicating that each subject’s alternative PRL offset was away from their highest-cone-density region. Figure 6D shows a comparison between the average cone density in the sector that subjects moved toward when establishing their alternative PRL and the direction they moved away from (i.e., 180 degrees away from the direction they moved toward). The average cone density was 15 *±* 10% lower in the direction that subjects moved toward compared to the opposite direction (t(6) = −4.26, p = 0.005, two-tailed paired t-test). This consistent trend to move away from the highest-density region immediately outside the scotoma is surprising because it suggests that changes in oculomotor behavior in this simulated scotoma paradigm are not random and are not driven by sampling maximization.

**Figure 6:**
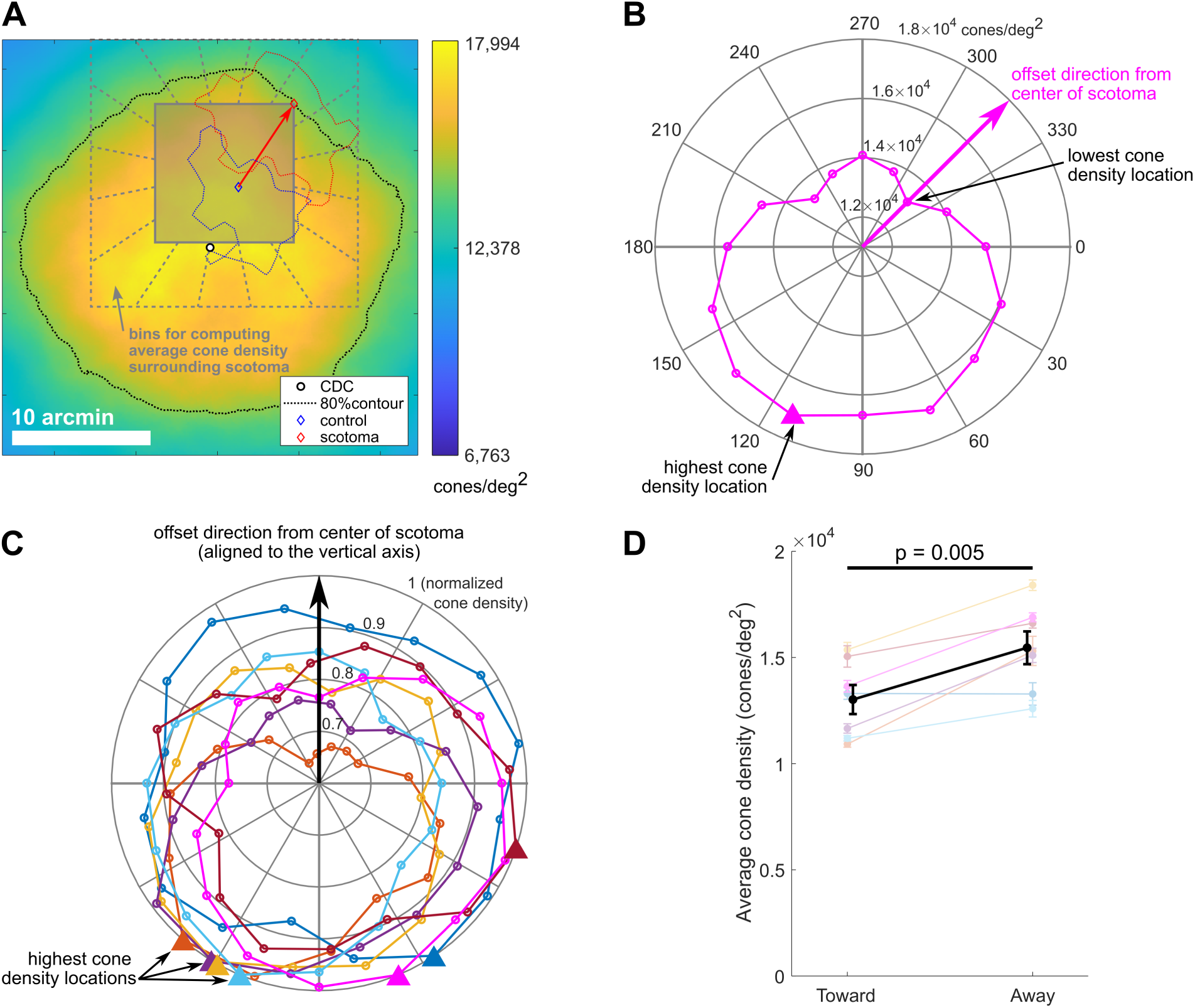
Alternative PRL offset angles relative to the cone density surrounding the scotoma. (A) The locations of the PRL in the control condition (blue diamond) and the alternative PRL in the scotoma condition (red diamond) are plotted on the cone density map for an example subject, with angular cone density values shown by the colorbar. The red arrow denotes the offset of the alternative PRL relative to the control condition. This example subject established an alternative PRL near the corner of the scotoma and closer to the edge of the high cone density region. The dashed gray lines show the bins used for calculating the average cone density surrounding the scotoma. (B) The average cone density (with units of cones per degree squared) in a 5-arcmin wide strip around the scotoma is plotted as a function of angle for this example subject. The offset direction is toward the lowest cone density location and away from the highest cone density location. (C) All seven subjects moved away from their highest cone density location. The black arrow denotes the offset direction of the alternative PRL, which was aligned to the vertical axis to enable comparison across subjects. The highest cone density locations are below the horizontal axis for all seven subjects. The units are normalized cone density. (D) The average cone density in the direction that subjects moved toward when establishing an alternative PRL was lower compared to the direction they moved away from (t(6) = −4.26, p = 0.005, two-tailed paired t-test; error bars denote SEM). This suggests that maximizing sampling density was not the primary factor influencing the formation of an alternative PRL in this task.

We further investigated the relationship between retinal anatomy and the alternative PRL offset direction to determine what may be influencing the change in oculomotor behavior in the scotoma condition. We observed that the offset of each subject’s alternative PRL from the center of the high-density region (i.e., the CDC) was in the same general direction as their control PRL was offset from the CDC (Figure 7A). Interestingly, in three of these subjects, the CDC location was not occluded by the scotoma, but even these subjects moved further from the CDC when establishing their alternative PRL instead of moving the stimulus toward the accessible CDC location.

**Figure 7:**
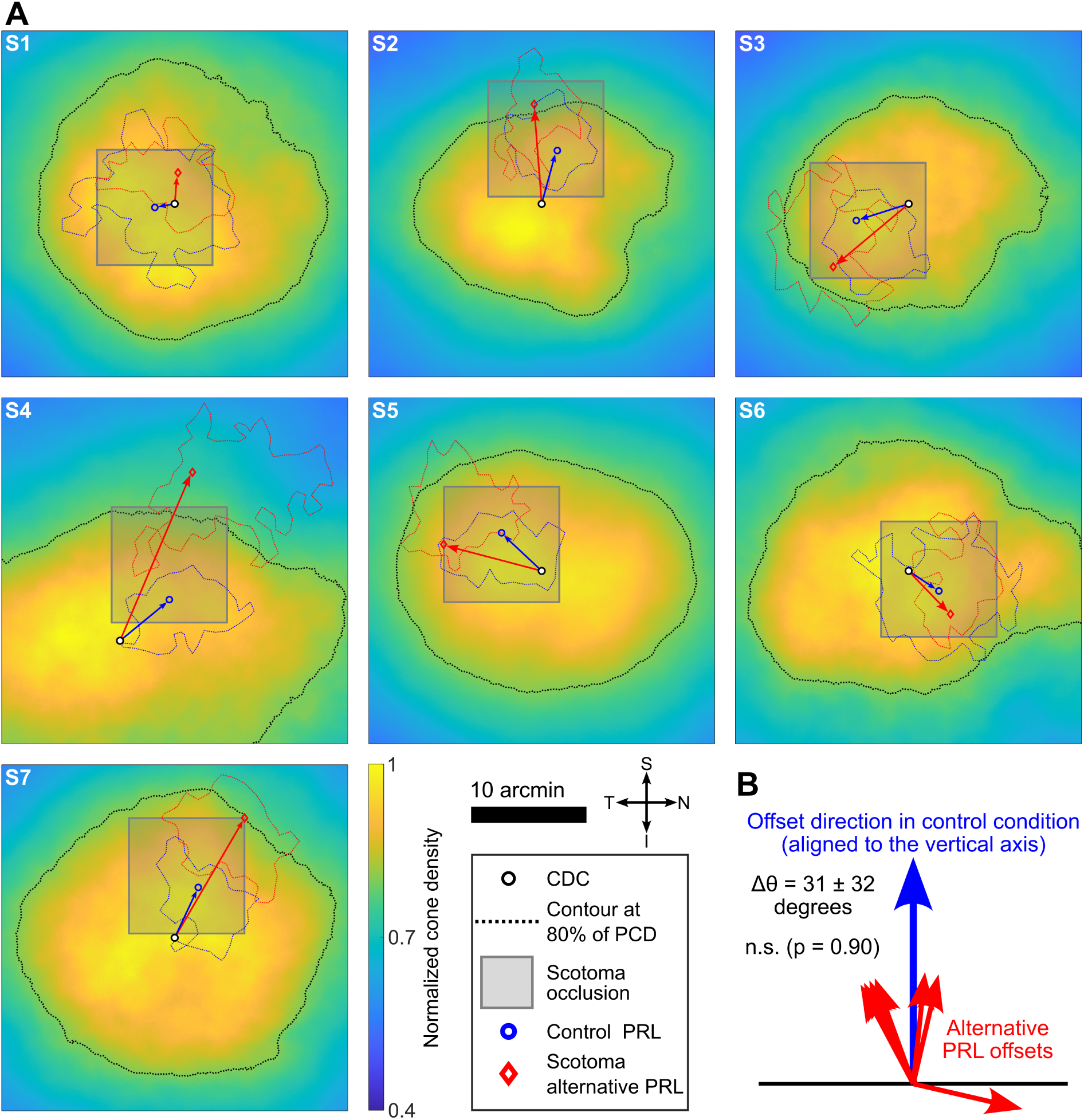
Alternative PRL offsets compared with the offset of the control PRL from the CDC. (A) The alternative PRL in the scotoma condition (red diamond) is offset in the same general direction from the CDC (black and white circle) as the control PRL (blue circle) across the 7 subjects. These locations are plotted on top of the normalized cone density map for each subject. The cardinal direction indicator in the legend refers to retinal coordinates (i.e., S refers to superior retina). (B) The difference in angle between the control (blue) and scotoma (red) offset vectors was 31 degrees on average, and there was no significant difference in the offset angles between the control and scotoma conditions (F(1,12) = 0.02, p = 0.90, Watson-Williams test for circular statistics). These results demonstrate that the alternative PRL is shifted further from the high-density region and in the same direction as the control PRL offset.

We compared the offset angles from the CDC location in both the control and scotoma conditions by aligning each subject’s control PRL offset angle (blue arrows) to the vertical axis, as shown in Figure 7B. The average magnitude of the difference in angle was 31 *±* 32 degrees, and there was no significant difference in the offset angles from the CDC location between the two conditions (F(1,12) = 0.02, p = 0.90, Watson-Williams test for circular statistics). Six of the seven subjects had angle offset magnitudes less than 30 degrees, while S1 had an angle offset of 103 degrees. This subject also had the smallest offset between their CDC and their control PRL, so we expect this small initial PRL offset is what led to a larger difference in angle compared to all other subjects. Rather than moving the stimulus toward the highest-density region of the retina to maximize sampling in response to the scotoma, the visuomotor system seems to set the alternate PRL further from the CDC in the same direction that the control PRL is already offset.

## Discussion

In this study, we investigated the impact of a simulated subfoveal scotoma in observers with normal vision. To the best of our knowledge, this is the first assessment of central vision that utilized retinal-contingent stimulation to precisely render a simulated subfoveal scotoma. We found that performance in a high-acuity discrimination task was significantly impaired for a 10-arcmin-wide scotoma, and observers changed their behavior in response to the scotoma by adopting an alternative PRL that was on average 5.5 arcmin—approximately half the scotoma width—away from their control PRL. There was not a systematic change in drift behavior across subjects in response to the scotoma, with the average observer maintaining a consistent directional bias and drift diffusion constant between the two conditions. The average offset direction of the alternative PRL was not toward the highest cone density region of the retina. Instead, the alternative PRL was offset in the same direction as the control PRL from the CDC, suggesting that subjects responded to the scotoma by moving the stimulus further from the region of higher cone density in the direction that they were already offset.

To establish an alternative PRL, we hypothesize that observers used microsaccades during the self-paced pause before each trial to offset their PRL from the location where the stimulus would appear. Since there were no systematic changes in drift behavior between the two conditions, microsaccades are likely responsible for the observed PRL offsets. In our data analysis pipeline, we only considered the video frames that included the stimulus, so we were unable to quantify eye motion before the stimulus onset. More research is needed to provide evidence for the role of microsaccades in establishing an alternative PRL in response to a simulated subfoveal scotoma.

In a control experiment with a subset of observers, we found that a 7-arcmin-wide scotoma did not elicit the same effect: observers did not establish an alternative PRL, and performance was not significantly impaired across subjects for this smaller scotoma. We hypothesize that there may be a minimum occlusion area before sizable changes in performance and behavior are exhibited. It is likely that an individual’s magnitude of fixational instability is closely related to the minimum scotoma size necessary to produce an effect. If a subject moved their eyes more, then they could potentially tolerate a larger scotoma before noticeable changes in performance or behavior would be observed. While our study did not systematically address this hypothesis about the minimum scotoma size, it laid the groundwork for future studies to investigate this effect.

Our findings rule out maximizing sampling density as a primary factor guiding the establishment of an alternative PRL in this task. Why might subjects use an alternative PRL with lower cone density rather than conveniently pick a retinal location with higher density? Recalling that subjects had a directional bias of drift that tended to bring the stimulus toward the CDC in both the control and the scotoma conditions (shown in Figure 4), we hypothesize that the offset of the alternative PRL is selected to maintain the sampling strategy that individual observers are accustomed to. In the control condition, observers tended to center a high-acuity stimulus slightly offset from the CDC location, and then move the stimulus toward the CDC location with their directional bias of drift over the course of a given trial. While there was variability across individual drift segments and observers, all subjects showed a tendency to drift in a specific direction. If the visuomotor system has developed this sampling strategy of moving a high-acuity stimulus toward higher density regions of the retina from a given starting location at the PRL, then it may be easier to maintain the same offset direction and directional bias of drift in the scotoma condition rather than switching to a different offset direction and/or dominant drift direction for the alternative PRL. While there are likely many motor and perceptual factors influencing behavior at this fine spatial scale, our results indicate that maximizing sampling density is not a primary factor influencing oculomotor behavior in this high-acuity task with a simulated scotoma. Interestingly, patients with central scotomas do not tend to select an alternative PRL with either the best visual acuity or contrast sensitivity of the region surrounding the scotoma (Rees et al., 2005, Bernard and Chung, 2018). Our results share some qualitative features with these prior observations by showing that higher cone density locations within the foveola do not coincide with a subject’s alternative PRL location in the simulated scotoma experiment.

Our study shows that subjects established an alternative PRL within the first block of scotoma trials. In subsequent scotoma blocks, subjects consistently offset in the same direction. This finding parallels studies with larger scotomas, where a stable alternative PRL was established rapidly in response to a simulated scotoma (Kwon et al., 2013, Liu and Kwon, 2016, Barraza-Bernal et al., 2018, Prahalad and Coates, 2020). For most subjects, we collected all scotoma blocks during a single experimental session, so we do not have sufficient evidence to establish the repeatability of the alternative PRL offset across multiple sessions. It remains an open question whether subjects would consistently adopt the same alternative PRL across days and weeks of repeated testing. Our finding that the alternative PRL is offset further from the CDC in the same direction as the control PRL would suggest, however, that subjects would likely return to the same general location on the retina, since the PRL has been shown to be very stable across multiple experimental sessions (Reiniger et al., 2021, Kilpeläinen et al., 2021).

In a control experiment conducted after the high-acuity task, each subject maintained fixation on a 5-arcmin-wide blinking square (3 Hz modulation frequency with 50% duty cycle) in both the presence and absence of the simulated scotoma for several 10-second viewing periods. Interestingly, we found no effect of the scotoma in the fixation experiment: the CDC-PRL offsets were 4.7 *±* 2.6 arcmin in the control condition and 5.5 *±* 2.4 arcmin in the scotoma condition (t(6) = −1.01, p = 0.35, two-tailed paired t-test). We posit that behavior did not change because the task did not require subjects to see the stimulus well, since there was no response required. On the contrary, subjects received immediate feedback if the stimulus drifted more than a few arcmin from the PRL, since the edges of the stimulus would become visible at the scotoma borders. In this way, subjects may have used the immediate feedback on stimulus position to recenter the blinking square behind the scotoma or along any of the scotoma edges without adopting a consistent offset. Results from this control experiment therefore raise questions about the influence of the task on oculomotor behavior with a subfoveal scotoma. Therefore, changes in fixation behavior may not be observed if the task does not require active discrimination of a target.

It would also be important to determine if and how the spatial frequency of the stimulus influences oculomotor behavior in the presence of a subfoveal scotoma. We kept the stimulus size constant in our study, with 5-arcmin-wide optotypes having 1-arcmin stroke widths (i.e., 30 cycles per degree for the fundamental spatial frequency). With adaptive optics correction, all subjects performed at ceiling for full-contrast stimuli of this size because the spatial frequency is around half of the sampling-limited spatial frequency of the retina, which exceeds 60 cycles per degree at the center of the fovea for most observers with normal vision (Curcio et al., 1990, Williams, 1988, Rossi and Roorda, 2010, Wang et al., 2019). If the stimulus spatial frequency was matched to an observer’s retinal sampling limit, developing an alternative PRL that was offset toward a location of lower sampling would not be an effective strategy. Would the system be flexible enough to account for this? Furthermore, if subjects were trained to adopt an alternative PRL that was closer to their CDC location, would they exhibit better performance compared with a baseline experiment with no training? These questions, while not addressed by our study, provide further avenues to explore the relationship between foveal anatomy and visual perception.

By using a simulated scotoma to temporarily occlude a small patch of the retina in healthy observers with normal vision, we can gain insight into oculomotor changes that accompany localized losses of visual function within the foveola. This paradigm may enable new discoveries in the disease progression and management of retinal degenerative diseases. It is important to note that our simulated scotoma paradigm differs in several ways from the visual experience of patients with retinal degenerative diseases. First, there was an immediate and abrupt appearance of the scotoma in our experiments. On the first trial with the simulated scotoma, the occluded region was completely blocked from seeing the stimulus, since the opacity of the scotoma was 100%. In retinal degenerative diseases like AMD, the development and progression of a central scotoma is gradual, with changes occurring on the timescale of weeks, months, and years (Bressler, 2002, Keane et al., 2015, Stahl, 2020).

The shape of our simulated scotoma was limited to a rectangular array due to constraints of the stimulus delivery mechanism of our AOSLO. We therefore used a square scotoma with abrupt edges, which is not characteristic of real scotoma shapes. We chose to render the scotoma with the same appearance as the background and no visible borders to make the scotoma invisible to observers. However, the photoreceptors within the simulated scotoma region still received input from the AOSLO illumination. In the case of a central scotoma, the photoreceptors within the scotoma are no longer functional, so they are unable to initiate any visual perception within the affected part of the visual field. Despite the differences mentioned above between simulated and real scotomas, our study highlights the fine control of fixational eye movements and the rapid adaptation of the visual system to a small subfoveal scotoma. These findings may advance our understanding of changes in retinal anatomy and oculomotor behavior during disease onset and progression that could lead to earlier diagnosis and treatment of retinal degeneration, leading to better outcomes and quality of life for patients.

## Methods

### Participants

Eight adult observers with normal vision and refractive errors less than 4 diopters in magnitude participated in this study. Three of the participants were female and five were male. Seven of the observers completed all experiment conditions in the study, while one participant completed only the fixation and 7-arcmin simulated scotoma conditions. Four participants were coauthors of this study, and four participants were naive to the study. The mean participant age was 28 years, with an age range of 23–31 years. Three observers who completed the main experiment also participated in the 7-arcmin control experiment. Each subject completed a preliminary screening during which ocular biometry, refractive error, and visual acuity were assessed. All procedures adhered to the ethical standards of the Research Subjects Review Board (RSRB) at the University of Rochester, which approved this study. Written informed consent was obtained from each participant after a thorough explanation of the study procedures and a detailed review of the materials in the consent form. See the participant summary table in Supplemental Information for more details.

### Retinal imaging and stimulation

We used a custom Adaptive Optics Scanning Light Ophthalmoscope (AOSLO) to simultaneously acquire high-resolution images of the retina while presenting visual stimuli during the experiments. Detailed specifications of the AOSLO are presented in Moon et al. (2024). Imaging and stimulus delivery were conducted using 680 nm light and a 1-degree by 1-degree square field of view. The AOSLO acquires videos at a frame rate of 30 Hz with a spatial resolution of 512 x 512 pixels, with each pixel subtending 0.12 arcmin. The AOSLO is equipped with a 940-nm illumination beam and a custom Shack-Hartmann wavefront sensor to measure the aberrations of the eye. A deformable mirror (DM97-08, ALPAO, Montbonnot, France) with an aperture of 7.2 mm and 97 actuators is positioned at a plane conjugate to the eye pupil and performs wavefront correction to compensate for the aberrations of the eye. Compensating for the eye’s aberrations is necessary to achieve cellular resolution within the central fovea, and it also enables stimuli to be delivered to the retina with near-diffraction-limited image quality. Backscattered light from the retina was focused onto a sub-Airy-disk pinhole (0.54 Airy-disk diameter) by a focusing lens and was incident on a photomultiplier tube (H7422-40, Hamamatsu, Shizuoka Pref., Japan) for detection. Custom data acquisition electronics and software then transformed the point-scanning temporal signal from the AOSLO into a spatial image of the photoreceptor mosaic. High-resolution images of the cone mosaic for each participant are shown in Figure 1 in Supplemental Information.

To render stimuli, the 680-nm imaging light source was rapidly modulated during the raster scan pattern using an acousto-optic modulator (TEM-210-50-10-680-2FP-SM, Brimrose Corp., Sparks Glencoe, MD). Using this high-speed modulator, the brightness of each pixel in the raster scan (i.e., the imaging field of view of the AOSLO) could be adjusted to render monochromatic stimuli with varying contrast levels. Decrement stimuli were implemented, so the stimuli appeared darker than the bright red background of the AOSLO raster scan.

### Experimental procedures

Data were collected across multiple experimental sessions, with each session lasting approximately 1–2 hours, and subjects completing an average of three sessions. At the beginning of each session, the right eye of each subject was dilated with one drop each of 1% tropicamide and 2.5% phenylephrine ophthalmic solutions approximately 15 minutes before starting the retinal imaging process (all testing was conducted monocularly in the right eye). After achieving pupil dilation, with a measured pupil diameter of 7 mm or greater, subjects were aligned to the AOSLO using a 3-axis translation stage fitted with a dental impression bar and temple pads, which kept the subject’s head stationary during the experiment. The subject’s pupil was aligned to the AOSLO imaging beam by observing the centration of the pupil in the live video feed of the custom wavefront control software. Closed-loop adaptive optics correction was then enabled while subjects viewed the dim red light of the AOSLO raster with the laser set to its minimum power. After achieving stable adaptive optics correction, the laser power was increased to its nominal level and imaging of the photoreceptor mosaic commenced. The average laser power levels used in these experiments were 27.8 *±* 1.5 *µ*W for 680 nm and 143.9 *±* 35.2 *µ*W for 940 nm, corresponding to 8% of the ANSI limit for photochemical hazards (applies to 680 nm light) and 3% of the ANSI limit for thermal hazards (applies to both 680 and 940 nm) for an exposure duration of one hour (Laser Institute of America, 2022).

In the first experimental session, subjects completed an acuity task using a 4AFC paradigm with Snellen E optotypes. The stimulus progression was self-paced in all conditions, with observers initiating each trial with a button press. The stimulus was presented for 500 ms in each trial, and a 1-second recording of the retinal cone mosaic was captured, with the stimulus presentation interval occurring in the middle of the 1-second recording. The stimulus size was constant, with a width of 5 arcmin, corresponding to a stroke width of 1 arcmin for the Snellen E optotypes (i.e., the 20/20 line of the eye chart). Due to the adaptive optics correction of ocular aberrations, observers performed at ceiling in the discrimination task when full contrast was used. To find an observer’s contrast threshold at the fixed stimulus size, we used an adaptive procedure with three interleaved QUEST staircases (Watson and Pelli, 1983), with each staircase containing 15 trials. We then took the average contrast of all QUEST staircases that converged and used this contrast for subsequent trials. The average contrast used in the control condition was 8 *±* 2% across subjects. Subjects then completed at least two blocks of the control condition, with approximately 60 trials per block, which enabled us to verify that performance was near threshold and to estimate each observer’s PRL.

This subset of the control data was then processed offline to identify each subject’s PRL, which was defined as the mean stimulus position on the retina across the subset of control trials (see details below on video processing). The procedure for determining the PRL also produced a high-resolution retinal image on which the PRL coordinates were marked. During the second experimental session, a 10-arcmin wide scotoma was stabilized at the location of the PRL during some experiment blocks. To implement retinal stabilization of the scotoma, stabilized viewing was first enabled in the data acquisition software and a reference frame was manually selected from a video frame with low motion. The reference frame selection was repeated until the stabilized view of the retina remained stationary during sustained fixation periods lasting at least 10 seconds. Next, the PRL coordinates were mapped onto the reference frame using normalized cross correlation in MATLAB to register the prior high-resolution retinal image with the reference frame. The scotoma was then rendered at the PRL coordinates with retinal stabilization enabled, meaning that the scotoma moved within the extent of the AOSLO raster scan to maintain a constant position on the retina. All pixels within the scotoma region had the same pixel value as the background illumination of the AOSLO raster scan, so the scotoma was invisible to observers unless the scotoma occluded the stimulus. The scotoma was placed one layer above the stimulus in the display software to ensure that the scotoma would cover up the stimulus if observers centered the stimulus at their PRL.

During this second experimental session, observers completed another round of QUEST to determine their contrast thresholds in the scotoma condition, using the same procedures outlined above. The average contrast in the scotoma condition was 27 *±* 7% across subjects, an increase by a factor of 3.3 *±* 0.3 compared to the control condition. For two observers who participated in an initial control experiment with the same stimulus contrast in both conditions, performance for the control trials was 60% correct for subject 1 and 84% correct for subject 2, and the performance dropped to chance in the scotoma condition: 23% correct for subject 1 and 24% correct for subject 2. In both the control and scotoma conditions, the stimulus was rendered at the center of the AOSLO raster and was stationary in world coordinates (i.e., the stimulus appeared at a fixed location in the raster and therefore moved across the retina due to fixational eye movements), while the scotoma was stabilized on the retina at each observer’s PRL in the scotoma condition. The same Snellen 4AFC task was used for both conditions. For subsequent blocks and sessions, the two conditions were interleaved. Subjects completed 3.0 *±* 0.5 control blocks and 3.5 *±* 0.5 scotoma blocks on average over the course of data collection.

The stimulus contrast in the 7-arcmin-scotoma control experiment that four participants completed was different from the stimulus contrast used in the 10-arcmin-scotoma main experiment. Due to the smaller scotoma size, participants performed near threshold in both the presence and absence of the simulated scotoma for the same stimulus contrast. Therefore, the contrast was set for each participant and was maintained in both conditions for the 7-arcmin-scotoma experiment, unlike the 10-arcmin-scotoma main experiment where different stimulus contrasts were used.

### Video processing and eye movement analysis

Videos recorded during the experiment were first concatenated to combine all individual trial videos collected during a given experimental block. By concatenating the individual trials before registering the video frames, we minimized the number of independent reference frames for each subject. The concatenated videos were then processed using strip-based video registration algorithms to obtain high-resolution retinal images, stabilized videos, and eye movement data (Stevenson et al., 2010, Zhang et al., 2021). The stimulus and scotoma positions were previously encoded in the video frames with digital markers—small white crosses—that were added by the video acquisition software but were not visible to the participant. Once the video frames were registered, stimulus motion on the retina was measured by tracking the stimulus coordinates (i.e., the position of the digital marker for the stimulus) in each frame of the stabilized video. In the same way, the scotoma coordinates were tracked by extracting the position of the scotoma digital marker in each video frame.

All video frames containing the stimulus were manually checked by a single observer to ensure that the registration software properly compensated for retinal motion. Any frames with poor registration—where consecutive frames shifted by more than one cone diameter (approximately 0.5 arcmin) in the vicinity of the stimulus presentation—were excluded from further analysis. This manual process guaranteed precise localization of the stimulus position on the retina in each stimulus frame. Each trial contained 15 stimulus frames. Across all conditions, we dropped any trials that contained fewer than 10 good frames after this manual frame checking process. In the scotoma condition, we also required that the stabilized scotoma location was maintained within a radius of 1 arcmin from the most frequent scotoma position to ensure robust stabilization. All frames that did not meet this criterion were excluded from further analysis, and trials with fewer than 10 good frames after applying these criteria were dropped. For control trials, we obtained an average of 167 *±* 29 good trials per subject, with a rejection rate of 8 *±* 5%. For scotoma trials, we obtained an average of 145 *±* 49 good trials per subject, with a rejection rate of 29 *±* 25%. The higher trial rejection rate in the scotoma condition was due to the additional scotoma stabilization requirement.

We then combined the stimulus coordinate data across multiple experiment blocks for a given condition by registering each block reference frame with a preselected high-resolution cone image using normalized cross correlation, similar to methods described in Kilpeläinen et al. (2021). This process enabled us to map all the stimulus coordinates onto a single coordinate system defined by a given subject’s high-resolution retinal image. We then used the stimulus coordinates on the retina to identify the PRL in the control condition and the alternative PRL in the scotoma condition. For each subject and condition, we generated a two-dimensional histogram of the stimulus distribution on the retina, using 1-arcmin bin widths and probability normalization (i.e., the sum of all bin contents was 1). For the analysis of the control PRL and the alternative PRL in the scotoma condition, we chose to consider the subset of trials with correct responses. The histograms contained 1,463 *±* 499 points on average, with each point representing a single measurement of the stimulus coordinates, which were acquired at the 30 Hz frame rate of the AOSLO. There was no significant difference in the number of points between the control and scotoma conditions (t(6) = 1.62, p = 0.16, two-tailed paired t-test). The retinal coordinates for the control and alternative PRLs were determined by finding the peak of the stimulus distribution histograms, corresponding to the retinal location with the highest probability of stimulation during each trial. All histograms yielded a single peak (i.e., there was one histogram bin with higher probability than all other bins for each subject and condition).

To generate contours containing the top 68% of data in each condition, we first sorted the histogram data by probability and then computed the cumulative sum from highest to lowest probability. We then selected a probability threshold corresponding to a cumulative sum of 0.68, or the top 68% of all stimulus locations. We then identified the contour corresponding to this probability threshold, which we plotted on top of the two-dimensional histogram to verify that the contour encompassed the high-probability region of the stimulus distribution map. We also quantified the percent overlap between the control and scotoma contours by finding the area of the intersection of the two 68% contours, dividing the intersection area by the total area of the control contour, and then multiplying by 100. In a comparison of the 7-arcmin-scotoma and 10-arcmin-scotoma experiments, we found that the average overlap was 66 *±* 16% for the 7-arcmin-scotoma experiment compared with 38 *±* 35% in the 10-arcmin results, but this trend was not significant across subjects (t(9) = −1.44, p = 0.185, unpaired t-test). It is likely that a balanced design with the same number of subjects in both the 10-arcmin and 7-arcmin experiments would yield a significant result for this comparison.

The high-resolution images of the retina were segmented using a semi-automated procedure to identify the cone centers. Custom ConeMapper software (Gutnikov et al., 2021) was used to identify cone coordinates by means of a fully convolutional network (Hamwood et al., 2019), which was trained on high-resolution AOSLO images of the human retina. The cone coordinates were then checked and manually corrected where necessary by a single trained observer. To compute cone density across the high-resolution retinal images, a Voronoi map was generated from the cone coordinates for each image, and the average area of the 150 nearest neighbors was calculated for each pixel in the image, following methods similar to Reiniger et al. (2021). The average area measurement was then inverted to compute the average cone density across the retinal image. The Cone Density Centroid (CDC) was found by taking the weighted centroid of the top 20% of cone density values within the cone density matrix. A contour was placed at 80% of the peak cone density value to show the retinal region with the highest cone density (Figures 4A, 6, and 7A).

In addition to the stimulus coordinates, which were extracted from the stabilized videos and used to determine the PRL, we also analyzed the eyetraces returned by REMMIDE, the software that we used for registration of the video frames (Zhang et al., 2021). We used 16 strips per frame for the strip-based registration, resulting in an effective sampling rate of 480 Hz for the eye movement data. Using the known stimulus coordinates extracted using the methods described above, we aligned the eyetrace data to the high-resolution cone image so that the x- and y-traces represented the stimulus motion on the retina. We excluded all eyetrace segments corresponding to video frames with poor stabilization, which were manually identified following the methods described above. We then segmented the eyetraces to identify periods of drift. We conducted additional analysis on these drift segments to measure the instantaneous drift velocity, amplitude, span, curvature, and diffusion constant.

Across subjects, we found no significant difference in drift amplitude (5.0 *±* 1.1 arcmin for control, 4.4 *±* 0.4 arcmin for scotoma, t(6) = 1.40, p = 0.21), span (2.8 *±* 0.6 arcmin for control, 2.9 *±* 0.6 arcmin for scotoma, t(6) = −0.26, p = 0.80), average speed (23.9 *±* 5.8 arcmin/s for control, 25.9 *±* 4.4 arcmin/s for scotoma, t(6) = −0.84, p = 0.43), curvature (10.4 *±* 2.6 arcmin*^−^*^1^ for control, 9.6 *±* 1.9 arcmin*^−^*^1^ for scotoma, t(6) = 0.87, p = 0.42), and diffusion constant (Figure 4E) (20.4 *±* 5.7 arcmin^2^/s for control, 21.7 *±* 5.9 arcmin^2^/s for scotoma, t(6) = −0.38, p = 0.72) between the two conditions (two-tailed paired t-tests).

To quantify the directionality of drift, we plotted a two-dimensional histogram of the instantaneous drift velocity across all drift segments for a given subject and condition. We then found the mean amplitude and angle of drift. This allowed us to compare the mean drift direction between the control and scotoma conditions, as shown in Figure 4D. To compute the probability that drift moved in the direction of the CDC, we found the angle between each subject’s PRL and CDC. We then divided the drift velocity histogram into two halves with a line through the origin that was perpendicular to the PRL-CDC angle. In other words, we considered all angles within *±* 90 degrees of the PRL-CDC angle to be toward the direction of the CDC, and all angles in the other half of the circle to be away from the direction of the CDC. We then summed over all histogram bins in each half to compute the probability of moving toward vs. away from the CDC location (Figure 4C). We also examined the mean stimulus position over the first 250 ms and the last 250 ms of each stimulus presentation interval to quantify changes in retinal stimulation over the course of the stimulation period.

Observers demonstrated a directional bias of drift, with the average drift direction tending to bring the stimulus toward the CDC. The mean stimulus position was offset from the CDC by 4.4 *±* 1.6 arcmin in the first 250 ms of the control trials, and offset by 3.8 *±* 1.6 arcmin in the second 250 ms (t(6) = 2.46, p = 0.049, two-tailed paired t-test). In the scotoma condition, the offsets were 7.4 *±* 3.9 arcmin in the first 250 ms and 6.5 *±* 3.6 arcmin in the second 250 ms (t(6) = 3.42, p = 0.014, two-tailed paired t-test). See Figure 2 in Supplemental Information for a comparison of these starting and ending offsets in both the control and scotoma conditions. The magnitude of the difference in mean drift velocity angles between the two conditions was 20 *±* 15 degrees on average (Figure 4D), with no significant difference in the mean drift direction between conditions (F(1,12) = 0.28, p = 0.61, Watson-Williams test for circular statistics).

During fixation, subjects often perform microsaccades. In our experimental conditions, 9 *±* 5% of trials contained at least one microsaccade. To verify that these microsaccades were not altering behavior between the two conditions, we computed the isoline contour area that encompassed the top 68% of all stimulus locations on the retina, including all the microsaccade trials that were excluded from the earlier drift analysis. We found no significant difference in the 68% isoline contour area between the two conditions (63 *±* 29 arcmin^2^ for control, 68 *±* 23 arcmin^2^ for scotoma, t(6) = 0.26, p = 0.81, two-tailed paired t-test). In summary, all metrics that we used to quantify drift behavior showed no significant effect of the scotoma.

We analyzed the angular distribution of the average cone density in the region surrounding the stabilized scotoma by considering a 5-arcmin-wide strip immediately outside the scotoma borders. We divided this strip into 16 angular bins, with each bin centered on a multiple of 22.5 degrees (i.e., 0, 22.5, 45, etc. degrees for the bin centers). We then computed the average cone density within each of these bins. Figure 3 in Supplemental Information shows the average normalized cone density surrounding the scotoma for each of the seven participants who completed this task (this figure has the same information as Figure 6C, but in retinal coordinates for each participant instead of aligned to a common orientation). We measured the angle between the center of the scotoma and the alternative PRL location, which we referred to as the direction that subjects moved toward when establishing their alternative PRL. We then found the bin center that was closest to this angle and used the corresponding average cone density value when determining the cone density in the direction that subjects moved toward vs. the direction they moved away from. The direction that subjects moved away from was exactly 180 degrees away from the direction they moved toward. To compare results of the angular distribution of cone density across subjects, we aligned all of the alternative PRL offset angles (i.e., the direction that subjects moved toward when establishing their alternative PRL) to the vertical axis (Figure 6C).

To compute the average difference in angle between two conditions (i.e., Δ*θ* in Figures 4D and 7B), we calculated the mean of the absolute value of the difference in angle between the two conditions. The standard deviation was also computed after finding the absolute value of the difference. By taking the absolute value, we can gain insight into the magnitude of the angular difference between the conditions, whereas the mean value of the difference is expected to be close to zero, since both positive and negative angles are returned after taking the difference between the two sets of angular data.

### Statistical analysis

We performed all statistical analysis using custom MATLAB code. The significance level was set at *α* = 0.05 for all statistical tests. For values reported in the text, we followed the format of (mean) *±* (standard deviation). All error bars shown in the figures correspond to the Standard Error of the Mean (SEM), except for Figure 2B, which has standard deviation error bars. For statistical tests of significance between the control and scotoma conditions, we used two-tailed paired t-tests for most comparisons, such as results shown in Figures 2, 3, 4, and 6. For the comparison in Figure 4C, we used a repeated-measures ANOVA test with Tukey’s Honestly Significant Difference (HSD) test. This statistical test allowed us to compare both the probabilities (toward vs. away, which was highly significant) and the conditions (control vs. scotoma, which was not significant). For comparing the offset magnitudes in the 7-arcmin and 10-arcmin conditions, we used an unpaired t-test because not all subjects completed both conditions. For circular statistics, such as those reported in Figures 4D and 7B, we used the CircStat MATLAB toolbox (Berens, 2009). We checked for significance between two sets of angles (corresponding to the two conditions) using the Watson-Williams test implemented in CircStat (Watson and Williams, 1956, Berens, 2009).

