## SupplementalInfo for "Subfoveal scotomas trigger fine-scale fixation reorganization: insights from retinal imaging and retinal-contingent stimulation"

### Supplemental Information

#### Participant summary

| Subject ID | Age (years) | Sex | Refractive error (diopters) | Axial length (mm) | PCD (cones/deg$^2$) | CDC-PRL offset (arcmin) |
|---|---|---|---|---|---|---|
| S1 | 29 | M | 0 | 23.03 | 15,739 | 2.7 |
| S2 | 29 | F | -0.25 | 23.02 | 17,280 | 7.1 |
| S3 | 28 | M | -1 | 24.26 | 20,897 | 1.7 |
| S4 | 29 | F | -3.5 | 24.32 | 16,153 | 5.5 |
| S5 | 26 | M | +0.5 | 23.78 | 13,847 | 4.8 |
| S6 | 31 | M | -3.5 | N/A | 18,958 | 2.9 |
| S7 | 29 | M | +0.75 | 24.75 | 17,994 | 6.7 |
| S8 | 23 | F | 0 | 23.77 | 15,470 | 7.8 |

#### Cone mosaic images

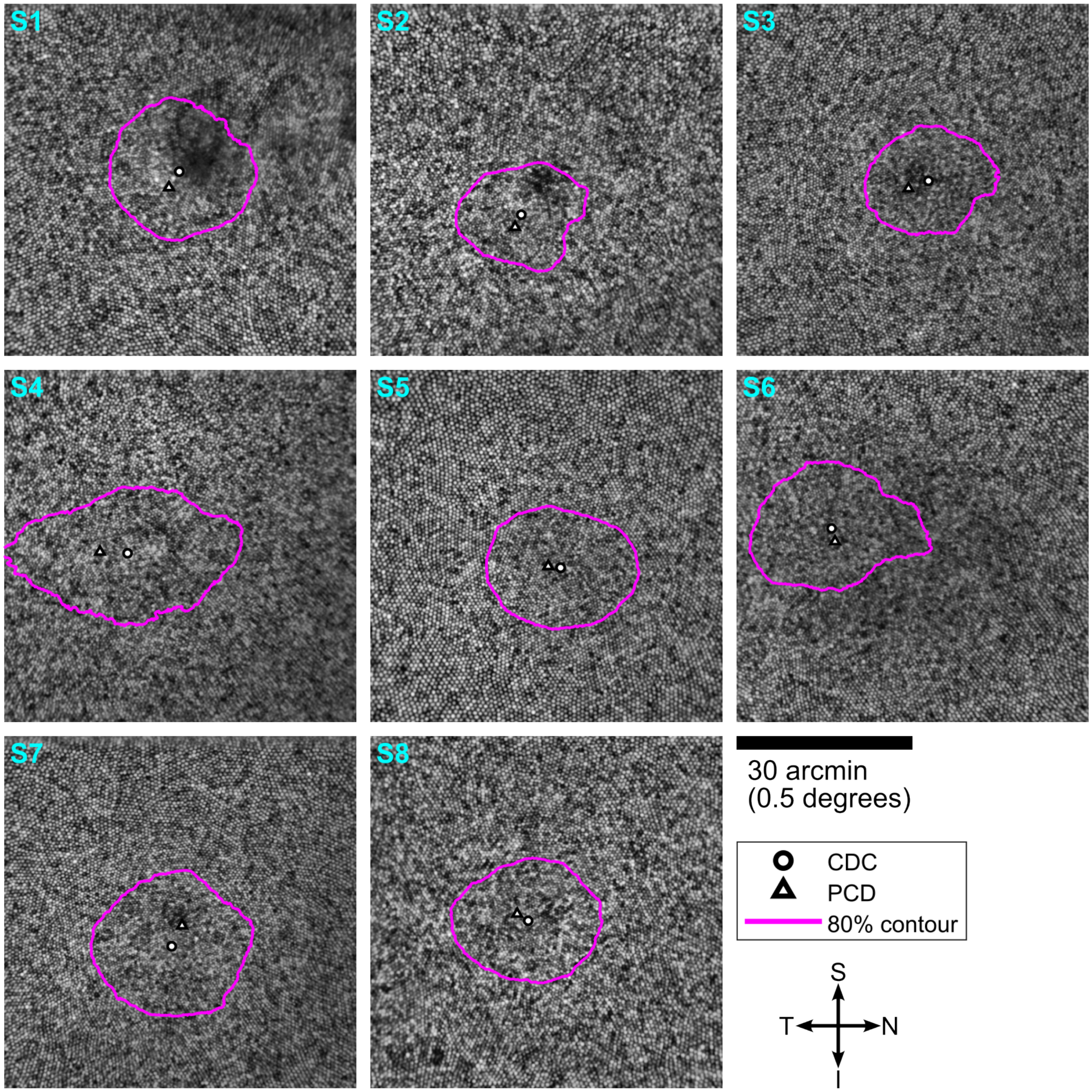


Figure 1: Images of the cone cone mosaic in the foveola for each participant. The Cone Density Centroid (CDC) and Peak Cone Density (PCD) locations are labeled. The magenta contour encloses the retinal area with cone density greater than 80% of the PCD value.

#### Offsets from CDC at start and end of trials

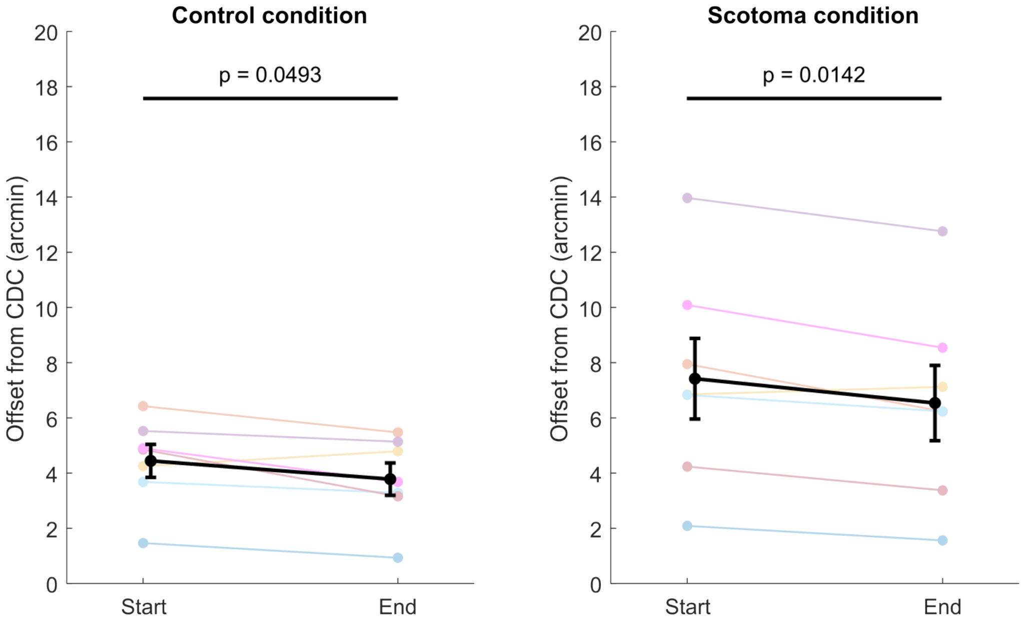


Figure 2: Offset magnitudes for the starting and ending segments of each trial, relative to the CDC location. The starting locations were computed using the first 250 ms of each stimulus presentation, and the ending locations used the last 250 ms of each stimulus presentation. Two-tailed paired t-tests were used for statistical analysis. Error bars denote standard error of the mean.

#### Angular distribution of normalized cone density surrounding the scotoma

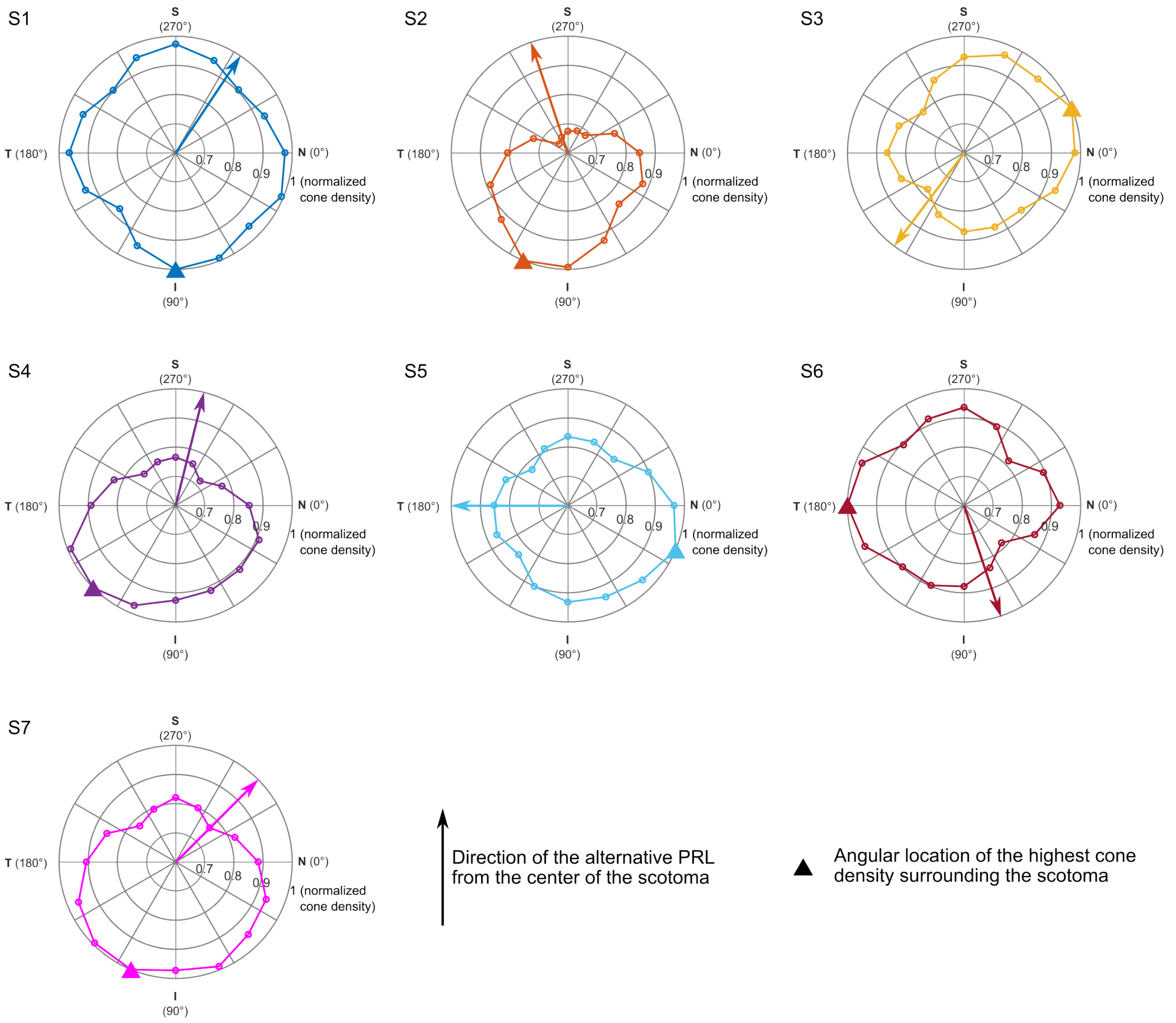


Figure 3: Normalized cone density in the 5-arcmin-wide strip surrounding the scotoma, plotted for individual subjects. The triangle denotes the angular location of the highest cone density surrounding the scotoma, and the arrow denotes the alternative PRL offset direction relative to the scotoma center.
